# A targeted, low-throughput compound screen in a *Drosophila* model of neurofibromatosis type 1 identifies simvastatin and BMS-204352 as potential therapies for autism spectrum disorder (ASD)

**DOI:** 10.1101/2022.11.11.516139

**Authors:** Alex Dyson, Megan Ryan, Shruti Garg, D. Gareth Evans, Richard A. Baines

## Abstract

Autism spectrum disorder (ASD) is a common neurodevelopmental condition for which there are no pharmacological therapies that effectively target its core symptomatology. Animal models of syndromic forms of ASD, such as neurofibromatosis type 1, may be of use in screening for such treatments. *Drosophila* larvae lacking *Nf1* expression exhibit tactile hypersensitivity following mechanical stimulation, proposed to mirror the sensory sensitivity issues comprising part of the ASD diagnostic criteria. Such behaviour is associated with synaptic dysfunction at the neuromuscular junction (NMJ). Both phenotypes may thus provide tractable outputs with which to screen for potential ASD therapies. In this study, we demonstrate that, while loss of *Nf1* expression within the embryo is sufficient to impair NMJ synaptic transmission in the larva, constitutive *Nf1* knockdown is required to induce tactile hypersensitivity, suggesting that a compound must be administered throughout development to rescue this behaviour. With such a feeding regime, we identify two compounds from a targeted, low-throughput screen that significantly and consistently reduce, but do not fully rescue, tactile hypersensitivity in *Nf1*^*P1*^ larvae. These are the HMG-CoA reductase inhibitor simvastatin, and the BK_Ca_ channel activator BMS-204352. At the NMJ, both compounds induce a significant reduction in the enhanced spontaneous transmission frequency of *Nf1*^*P1*^ larvae, though again not to the level of vehicle-treated controls. However, both compounds fully rescue the increased quantal size of *Nf1*^*P1*^ mutants, with simvastatin also fully rescuing their reduced quantal content. Thus, the further study of both compounds as potential ASD interventions is warranted.

**Significance Statement:** No therapies currently exist that consistently and effectively target the core symptoms of autism spectrum disorder (ASD), which include altered responses to sensory stimuli. Previously it was shown that *Drosophila* larvae lacking expression of ASD-associated *Nf1* display a heightened response to a mechanical stimulus and increased neuronal excitability, likely due to excessive Ras activity. Here, out of a screen for compounds targeting such mechanisms, we identified simvastatin and BMS-204352 to reduce the likelihood of a response in *Nf1*^−/-^ larvae following mechanical stimulation. These compounds also improved synaptic transmission defects at the neuromuscular junction. Such findings support the further study of these drugs as potential ASD therapies in the clinic.

## Introduction

Autism spectrum disorder (ASD) is a common neurodevelopmental condition affecting 1 – 2% of children (Baird et al., 2006; Maenner et al., 2020). Clinically, it is characterized by impairments in social communication alongside repetitive and restricted behavior and interests, which include altered responses to sensory stimuli (American Psychiatric Association, 2013). Such sensory impairments may directly contribute to other ASD symptoms, and correlate with worsening outcomes on several quality-of-life measures. Thus, they may provide an important target for therapeutic intervention (Lundqvist, 2015; Robertson and Baron-Cohen, 2017; Lin and Huang, 2019). However, despite the lifelong impact on an individual’s quality of life that ASD can impose (Lord et al., 2020), and the substantial economic burden arising from the need to support those affected (Buescher et al., 2014; Leigh and Du, 2015), there are currently no treatments that consistently and effectively improve abnormal sensory sensitivity or other aspects of core ASD symptomatology (Hyman et al., 2020).

While ∼75% of ASD cases are idiopathic, approximately 4 – 5% occur in association with a monogenic neurodevelopmental syndrome (Fernandez and Scherer, 2017). Because syndromic forms of ASD have a known single causative mutation, they are comparatively simpler and offer a more tractable model from a biomedical research perspective (Sztainberg and Zoghbi, 2016). One such condition is Neurofibromatosis Type 1, an autosomal dominant disorder arising from loss of function of the *NF1* gene on chromosome 17 (Gutmann et al., 2017). The prevalence of ASD amongst individuals with Neurofibromatosis Type 1 is estimated at 10 – 25%, with up to a further ∼20% exhibiting some form of clinically relevant symptomatology (Garg et al., 2013; Walsh et al., 2013; Adviento et al., 2014; Plasschaert et al., 2015; Morris et al., 2016; Eijk et al., 2018). Furthermore, it is becoming increasingly apparent that ASD and Neurofibromatosis Type 1 share an overlapping pathophysiology. Thus, animal models of Neurofibromatosis Type 1 possess significant potential in screening for novel therapies for ASD (Molosh and Shekhar, 2018; Kaczorowski et al., 2020).

An effective drug screen foremost requires a suitable model of the disorder. Target-based, *in vitro* approaches have not proven suitable for conditions like ASD in which the underlying disease biochemistry is poorly understood and likely involves the dysfunction of multiple pathways. They also do not routinely permit behavioral outputs as measures of drug efficacy (Pandey and Nichols, 2011; Strange, 2016). More traditional, pre-clinical model organisms, such as mice are equally unfeasible, given their high cost of maintenance, long generation time, and relatively small litters (Bell et al., 2009). By contrast, the fruit fly, *Drosophila melanogaster*, overcomes many of these limitations and provides a useful bridge between the two model systems (Bell et al., 2009; Pandey and Nichols, 2011; Strange, 2016). The potential of *Drosophila* in neurodevelopmental drug discovery was exemplified by a screen of 2,000 small molecules for their ability to rescue glutamate-induced lethality in a fly model of ASD-associated Fragile X syndrome (Chang et al., 2008). Indeed, identification of GABA as a hit compound lent support for the use of GABA-promoting drugs in clinical trials (Braat and Kooy, 2015), although the success of these has been mixed (Berry-Kravis et al., 2012; Erickson et al., 2013; Erickson et al., 2014; Ligsay et al., 2017; Veenstra-VanderWeele et al., 2017).

*Drosophila* express a highly conserved ortholog of *NF1* (The et al., 1997) with similar molecular, cellular, and behavioral functions to its mammalian counterpart (Guo et al., 2000; Walker et al., 2006). Accordingly, *Nf1*^*-/-*^ flies display phenotypes analogous to ASD in the clinic, including altered communication (Moscato et al., 2020), disrupted sleep (Bai and Sehgal, 2015), and repetitive behaviors in the form of excessive grooming (King et al., 2016; King et al., 2020). More recently, it was shown that *Drosophila* larvae lacking *Nf1* expression exhibit an increased likelihood of responding to a mechanical stimulus, thought to mirror the sensory sensitivity abnormalities comprising part of the ASD diagnostic criteria (Dyson et al., 2022). Here, we exploit this phenotype in a targeted, low-throughput screen to identify compounds that improve tactile hypersensitivity in *Nf1*^*P1*^ larvae. Any such leads may therefore have potential in managing ASD symptomatology in affected individuals. We start by investigating the temporal requirements for *Nf1*, in order to determine when compounds must be administered for optimal activity. Then, using a protocol of feeding drugs during both embryonic and larval stages, we identify two compounds, simvastatin and BMS-204352, as capable of consistently improving, but not fully rescuing, the enhanced response to mechanical touch. Finally, we demonstrate that both compounds also reduce *Nf1*-associated synaptic transmission deficits at the neuromuscular junction (NMJ).

## Materials and Methods

### Fly lines and maintenance

The *Nf1*^*P1*^ mutant and its parental K33 line used in this study (The et al., 1997) were initially obtained from Seth Tomchik (University of Iowa, USA), where they were both backcrossed into the *w*^*CS10*^ background such that K33 acts as an isogenic control, as described previously (King et al., 2016; Dyson et al., 2022). For the temperature-dependent knockdown of *Nf1* at different developmental stages, *elav*^*c155*^*-GAL4;tubP-GAL80*^*ts*^ was crossed to either *UAS-Nf1*^*RNAi*^*;UAS-Dicer2* (experiment) or *UAS-GFP*^*RNAi*^*;UAS-Dicer2* (control). These lines were generated by combining the following constructs together: *elav*^*c155*^*-GAL4* (Bloomington, #458) (Giachello and Baines, 2015), *tubP-GAL80*^*ts*^ (McGuire et al., 2004), *UAS-Nf1*^*RNAi*^ (VDRC ID #109637) (King et al., 2016), *UAS-GFP*^*RNAi*^ (Bloomington #9331) (Roignant et al., 2003), and *UAS-Dicer2* (Dietzl et al., 2007). In these experiments, animals were maintained at either 30 °C to permit GAL4-dependent expression of *UAS-Nf1*^*RNAi*^ and thus knockdown of endogenous *Nf1*, or at 18 °C to facilitate GAL80-dependent repression of GAL4, thereby allowing the expression of endogenous *Nf1* (McGuire et al., 2004). Progenitor flies were set up to mate and left to deposit embryos on grape agar plates, which were collected at 4-hour intervals before being transferred to food-containing vials. For the constitutive knockdown of *Nf1*, embryos and larvae were maintained constantly at 30 °C until the experiment. For the embryonic knockdown of *Nf1*, embryos were maintained at 30 °C for 17 – 21 h after egg laying (AEL), and then transferred to 18 °C (Ashburner et al., 2005). For the larval knockdown of *Nf1*, embryos were maintained at 18 °C for 42 – 46 h AEL, and then transferred to 30 °C (Ashburner et al., 2005). For all other experiments, lines were maintained at 25 °C. Fly food was standard cornmeal medium. All flies were kept in a 12:12 light:dark cycle.

### Compound administration

Compounds that were tested for their ability to improve tactile hypersensitivity in *Nf1*^*P1*^ larvae are listed in Table 1. All compounds were made up as stock solutions in DMSO and stirred into molten fly food cooled to ≤ 60 °C. Typically, progenitor flies were maintained on food containing either compound or vehicle (DMSO) for ≥ 3 days before being transferred onto fresh compound- or vehicle-containing food, from which larvae for the experiment were collected. However, for experiments examining embryonic exposure to compound, progenitor flies were maintained on food containing either compound or vehicle (DMSO) for ≥ 3 days before being transferred onto standard food (lacking compound), from which larvae for the experiment were collected. For experiments examining larval exposure to compound, progenitor flies were maintained on standard food before being transferred onto fresh compound- or vehicle-containing food, from which larvae for the experiment were collected. DMSO concentration was 0.05 % for all experiments presented in this study.

**Table 1.** List of compounds screened for their ability to improve tactile hypersensitivity in *Nf1*^*P1*^ larvae. Compounds were selected based on either their ability to inhibit Ras/MAPK activity or to modulate ion channel function, mostly with the expectation of reducing neuronal excitability. As part of the ‘Ras/MAPK’ targeting group, inhibitors of histone deacetylase (HDAC) and CDK4/6 were included since they have been demonstrated to improve viability in *Drosophila* models of other RASopathies (Das et al., 2021).

| Process targeted | Compound | Mechanism of Action | Source |
| --- | --- | --- | --- |
| Ras/MAPK signalling | Lovastatin | HMG CoA-reductase inhibitor | Generon |
|  | Fluvastatin Sodium |  | Generon |
|  | Simvastatin |  | Fluorochem |
|  | Atorvastatin Calcium |  | Generon |
|  | Binimetinib | MEK inhibitor | Generon |
|  | U0126 |  | Cambridge Bioscience |
|  | GDC-0973 (Cobimetinib) |  | Generon |
|  | PD 0325901 |  | Merck |
|  | Salirasib | Ras inhibitor | Generon |
|  | GNE 3511 | DLK inhibitor | Generon |
|  | Palbociclib HCl | CDK4/6 inhibitor | Generon |
|  | Vorinostat | HDAC inhibitor | Generon |
|  | Belinostat |  | Generon |
|  | Trichostatin A |  | Cambridge Bioscience |
|  | BAY293 | Ras/SOS1 interaction inhibitor | Generon |
| Ion channel function | ML 6733 | K<sub>2P</sub> activator | Generon |
|  | BMS204352 | BK<sub>Ca</sub> channel activator | Cambridge Bioscience/Scientific Laboratory Supplies |
|  | BAY K 8644 | L-type Ca<sub>v</sub> channel activator | Stratech Scientific Ltd |
|  | Lamotrigine | Sodium channel blocker | Fisher Scientific UK Ltd |
|  | SKA 31 | SK channel activator | Bio-Techne |

### Tactile sensitivity and compound screening

To assess tactile sensitivity, a mechanical stimulus was applied briefly to the posterior end of each wall-climbing third instar larva (of both sexes), as described previously (Dyson et al., 2022). Larvae were counted as ‘responding’ only if they exhibited a full, 360° rolling motion. All such experiments were carried out at room temperature by an experimenter blinded to genotype and/or treatment. For the initial screen, all compounds were tested at 50 µM on *Nf1*^*P1*^ larvae and compared against *Nf1*^*P1*^ larvae raised on an equivalent concentration of DMSO vehicle (*n* = 20 per treatment group). Data for each compound were accepted if vehicle-treated *Nf1*^*P1*^ larvae exhibited the expected significant increase in likelihood of nocifensive response compared to vehicle-treated K33 larvae (*n* = 20). For the compound screen, data were normalised with the equation: (no. of compound-treated responders/no. of vehicle-treated *Nf1*^*P1*^ responders) x 100, such that the number of *Nf1*^*P1*^ larvae responding to the stimulus = 100%. Compounds found to exert a significant effect in the initial screen were then tested in four independent validation trials (*n* = 20 per treatment group per trial) at 50 µM. For compound validation, % responding larvae per trial (i.e. (no. of responders/20) x 100) for each genotype was calculated.

### Larval Crawling

A 77 cm^2^ arena formed of 2 % agarose (depth ∼2 mm) within the lid of a clear 96-well plate was placed into a DanioVision observation chamber. A 2 – 3 mm moat was maintained between the agar and the edge of the lid, which was filled with 5 M NaCl to deter larvae from crawling off the agar. Wall-climbing third instar larvae (of both sexes) were rinsed in ddH_2_O, then placed onto the centre of the arena, and left to acclimatise for 30 s. EthoVision XT Video Tracking Software (part of DanioVision) was used to measure total distance travelled over a 3-minute period under white light. Experiments were carried out at room temperature by an experimenter blinded to genotype and/or treatment. Only larvae that remained on the agar for the entire recording period were included in our analyses.

### Electrophysiology

Wall-climbing third instar larvae (of both sexes) were fillet-dissected to permit the recording and analysis of excitatory junction potentials (EJPs) and miniature EJPs (mEJPs) in HL3 saline + 1.5 mM CaCl_2_ (Stewart et al. 1994), as described previously (Dyson et al., 2022). EJP amplitude and resting membrane potential were calculated using Clampfit 10.3 (Molecular Devices), while mEJP amplitude and frequency were calculated using MiniAnalysis (Synaptosoft Inc. GA, USA). Amplitudes were corrected using the equation v’ = E(ln[E/(E-v)]), where v’ refers to the corrected amplitude, v is the recorded amplitude, and E is the driving force, assumed to be equal to resting membrane potential if the reversal potential is 0 mV (Feeney et al., 1998). Quantal content was calculated as corrected EJP amplitude/corrected mEJP amplitude. All recordings were carried out blind to genotype or treatment.

### Statistical Analysis

All statistical analyses were carried out using GraphPad Prism (version 8). Pairs of quantitative data sets were compared via a two-tailed, unpaired student’s t-test, while ≥3 quantitative data sets were analysed via either one-way (ungrouped) or two-way (grouped) ANOVA, followed by Tukey’s post-hoc test. Fisher’s exact test was used to compare two sets of categorical variables (e.g. vehicle-vs. compound-treated *NF1*^*P1*^ larvae). All analysis was carried out on raw data prior to normalisation. Statistically significant p values are given in the figures, while non-significant p values that are nevertheless relevant to interpreting the data are given in the figure legend.

## Results

### Constitutive knockdown of *Nf1* is required to induce tactile hypersensitivity, while knockdown of *Nf1* in the embryo alone is sufficient to impair synaptic transmission

Because Neurofibromatosis Type 1 and ASD are primarily developmental disorders, it is probable that they arise, at least partially, from aberrations in brain development and/or function during early life stages that lead to permanent changes in postembryonic behaviour (Yenkoyan et al., 2017; Courchesne et al., 2019). These aberrations may be irreversible, and, consequently, less susceptible to clinical intervention in later life (de la Torre-Ubieta et al., 2016). To account for this, we first investigated when the NF1 protein is required during the *Drosophila* life cycle to regulate larval tactile sensitivity. We also focused attention to synaptic transmission at the NMJ, since deficits in the latter may be associated with the hypersensitivity phenotype (Dyson et al., 2022). We used the UAS/GAL4/GAL80^ts^ system to restrict *Nf1* expression to either embryonic or larval stages (McGuire et al., 2004). In these experiments, larvae in which *Nf1* was knocked down via RNA interference were compared to a control line in which a dsRNA construct against GFP was expressed instead.

As expected, constitutive knockdown of *Nf1* resulted in a significant increase in the likelihood of a larva exhibiting a nocifensive response following a brief, typically innocuous, mechanical stimulation (Figure 1A). However, this was not observed when *Nf1* knockdown was restricted solely to the embryonic period (Figure 1B) or, alternatively, to post-embryonic larval stages (Figure 1C). To completely rule out the possibility that some non-specific effect of raising larvae at 18 °C, independent of GAL80^ts^ expression, diminishes the nocifensive response, we also exposed *Nf1*^*P1*^ mutant larvae and the isogenic K33 controls to the same shifts in temperature. In all such controls, the number of *Nf1*^*P1*^ larvae displaying the nocifensive response was significantly greater than that of control larvae (Figure 1D-F). Thus, we conclude that the NF1 protein is involved in regulating normal tactile sensitivity during both embryogenesis and larval development.

**Figure 1.**
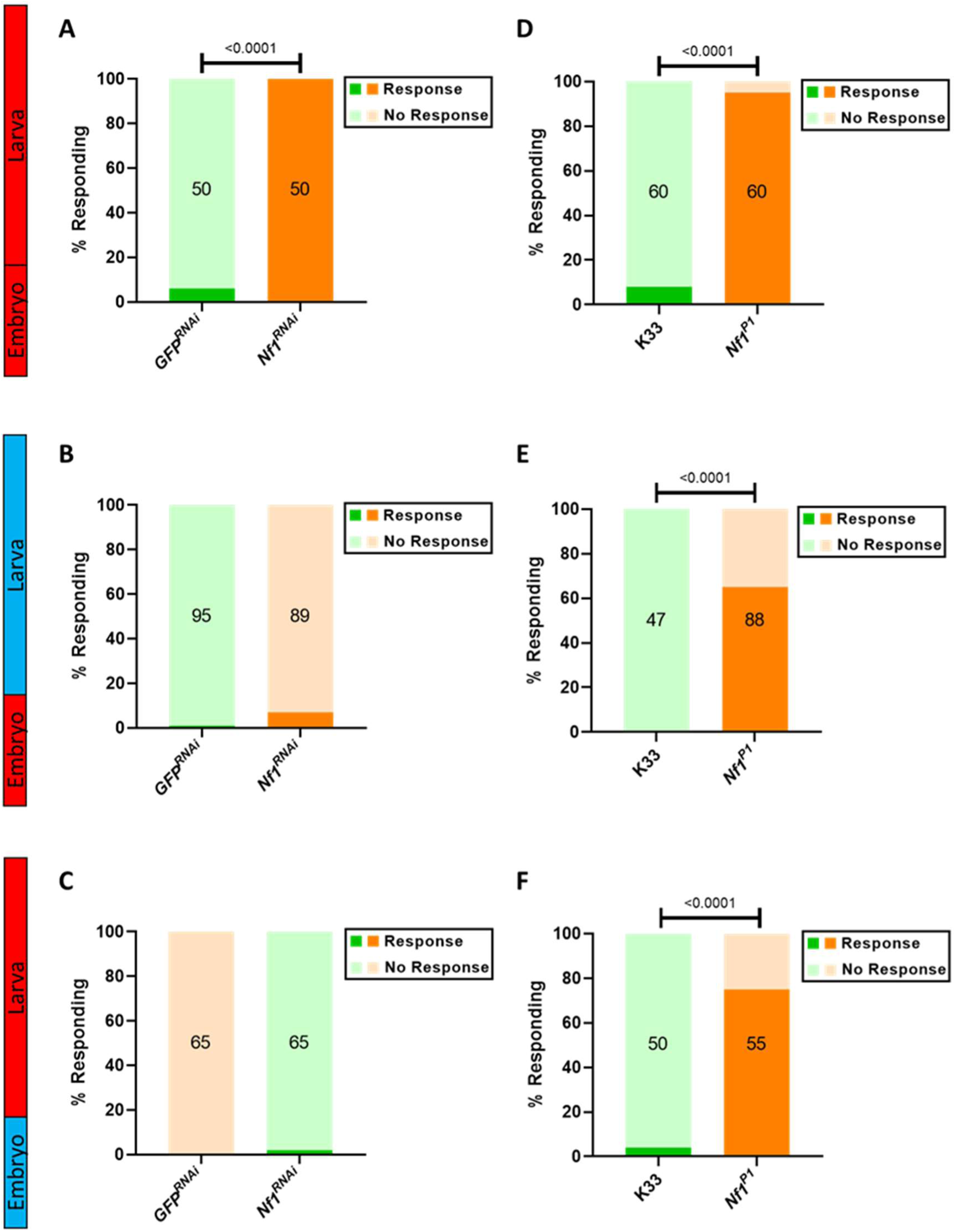
Constitutive knockdown of *NF1* is required to induce tactile hypersensitivity in third instar larvae. Abbreviated genotypes for the lines tested are given in the figure panels. *GFP*^*RNAi*^ (green) and *Nf1*^*RNAi*^ (orange) refer to lines in which GAL4 is expressed under the control of *elav* to drive expression of either UAS-*GFP*^*RNAi*^ or UAS*-Nf1*^*RNAi*^, respectively, and UAS-*Dicer2*, with GAL80^ts^ expressed under the control of the *tubulin* promoter. All transgenic constructs are hemi- or heterozygous in the larvae tested. *A)* Constitutive knockdown of *Nf1* expression (*Nf1*^*RNAi*^) throughout all life stages results in larval tactile hypersensitivity, as indicated by a significantly greater number of larvae responding to a mechanical stimulus compared to *GFP*^*RNAi*^. *B)* Knockdown of *Nf1* only within the embryo has no significant impact on the number of responding larvae (p=0.0580), nor does *C)* knockdown of *Nf1* within the larval stages (p>0.9999). *D-F)* K33 and *Nf1*^*P1*^ larvae were also subjected to the same shifts in temperature as those required for constitutive, embryonic, and larval knockdown of *Nf1* respectively. Regardless of the temperature paradigm, *Nf1*^*P1*^ larvae always demonstrated a significantly greater likelihood of displaying the nocifensive response following stimulation. Numbers within each bar represent the sample size for that group. For the ease of comparing groups in which different sample sizes were used, raw data have been presented as percentages within the figure. Statistical comparisons via Fisher’s exact test were nonetheless carried out on raw data prior to normalisation.

Next, we examined the effect of identical shifts in *Nf1* expression on synaptic transmission at the NMJ. In *Nf1*^*-/-*^ mutants, the frequency of spontaneous transmission events (mEJPs) is significantly increased, while evoked release (quantal content) is significantly reduced. This change is seemingly compensated for by an increase in postsynaptic input resistance, as evidenced via an enhanced quantal size (mEJP amplitude), rendering the amplitude of evoked events (EJPs) unchanged (Dyson et al., 2022). These changes are similarly observed following constitutive *Nf1* knockdown (Figure 2A-F), as well as when knockdown of *Nf1* is restricted to the embryonic period (Figure 2G-L). However, knockdown of *Nf1* during larval stages is insufficient to disrupt synaptic transmission (Figure 2M-R). Together, these data indicate an early developmental requirement for *Nf1* in regulating synaptic transmission at the larval NMJ, but that the role of *Nf1* in sensory sensitivity requires expression at both embryonic and larval stages.

**Figure 2.**
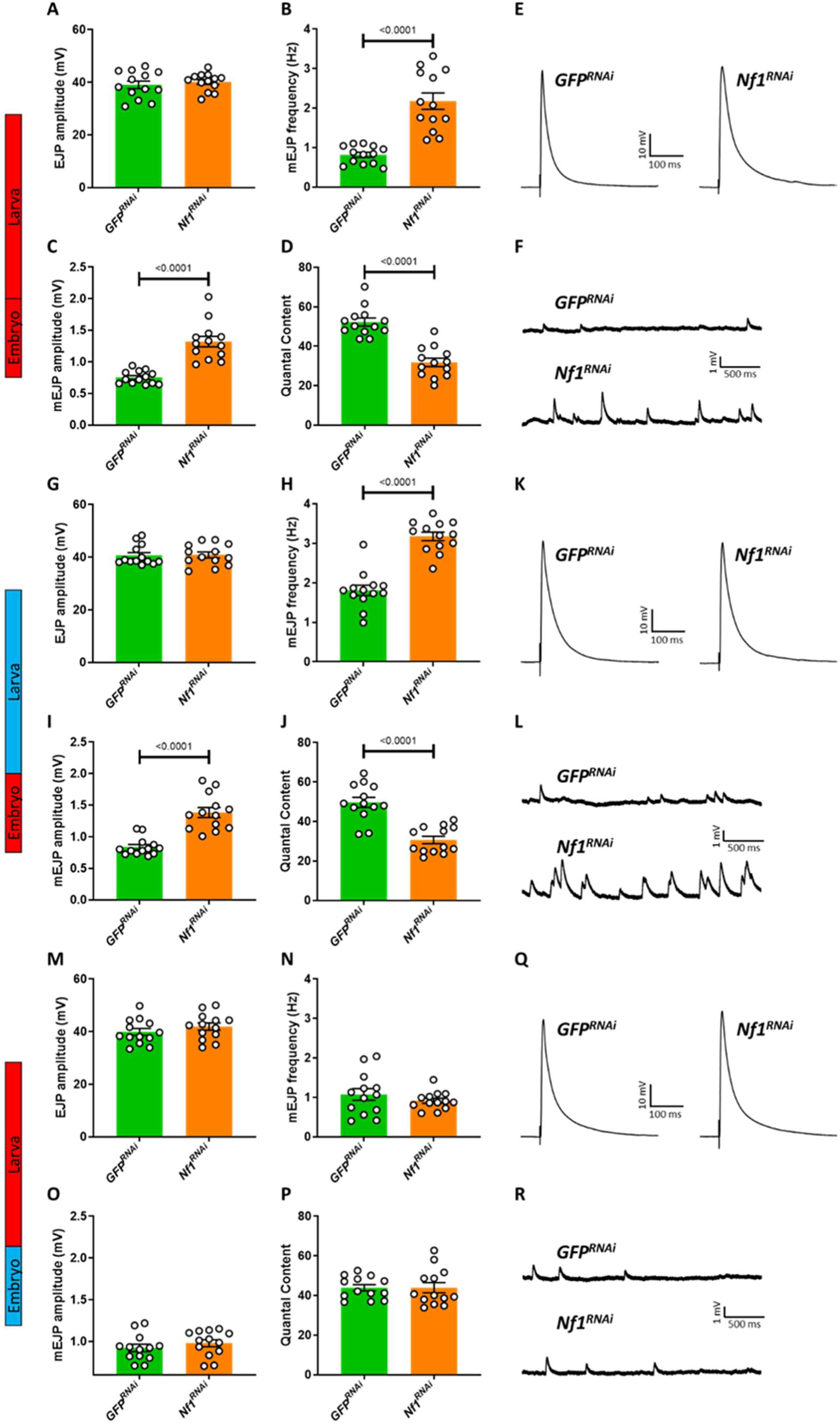
Knockdown of *NF1* in the embryo is sufficient to induce synaptic transmission deficits in the third instar larval stage. Full genotypes for those abbreviated in the figure (i.e. *GFP*^*RNAi*^ and *Nf1*^*RNAi*^) are explained in the legend for Figure 1. *A-F*) Raising *Nf1*^*RNAi*^ larvae at 30 °C, sufficient to ensure constitutive knockdown of *Nf1*, mimics the *Nf1*^*-/-*^ larval phenotype (Dyson et al., 2022) in that it does not affect EJP amplitude (p=0.5334) but increases mEJP frequency and amplitude, and reduces quantal content. *G-L)* Knockdown of *Nf1* within the embryonic period has a similar effect on synaptic transmission as constitutive knockdown on all parameters examined, including no significant impact on *G)* EJP amplitude (p=0.8967). *M-R)* In contrast, knockdown of *Nf1* only during larval stages has no effect on *M)* EJP amplitude (p=0.3010), *N)* mEJP frequency (p=0.3230), *O)* mEJP amplitude (p=0.3694) or *P)* quantal content (p=0.9998). Data in each panel were analysed via unpaired, two-tailed student’s *t*-test. For each experiment, *n =* 13. Data are presented as mean ± SEM.

### A targeted screen identifies two compounds sufficient to improve tactile hypersensitivity in *Nf1*^*P1*^ larvae

To expedite the screening process, we opted to assay compounds for their ability to improve tactile hypersensitivity, rather than aberrant synaptic transmission. As knockdown of *Nf1* in both the embryonic and larval periods is necessary to induce this behavioural phenotype, it is plausible that, for a compound to rescue such behaviour, it must likewise be present throughout both stages. To achieve this, progenitor flies were provided compound-containing food for ≥3 days before being transferred onto fresh food containing the same compound on which eggs would be laid and larvae would develop. This was to ensure that the compound was present both during the embryonic and larval stages, as feeding compounds to mated females results in significant transfer to embryos (Marley and Baines, 2011). Compounds were chosen for their ability to either diminish Ras/MAPK signaling, since knockdown of either *Ras85D* or *Ras64B* fully rescues both hypersensitivity and synaptic deficits in *Nf1*^*P1*^ larvae (Dyson et al., 2022), or for their ability to modulate ion channels, typically in favor of reducing neuronal excitability. All drugs were tested at 50 µM (i.e. the concentration added to fly food), although the concentration reaching the central nervous system (CNS) is unknown. While three of the compounds tested proved toxic at this concentration (Fluvastatin, PD0325901, and Trichostatin A), we identified two compounds that significantly reduced the number of *Nf1*^*P1*^ larvae responding to mechanical stimulation (Figure 3). These were the HMG CoA-reductase inhibitor simvastatin, and the big conductance calcium-activated potassium channel (BK_Ca_) activator BMS-204352. Four subsequent independent trials demonstrated that the effect of each compound was consistent, with a significantly lower mean percentage of *Nf1*^*P1*^ responders in the compound-treated vs. vehicle-treated groups (Figure 4A-B). It is worth noting, however, that each compound exerted a significant effect in only three out of four trials when comparing treatment groups within the trial itself (Table 2).

**Table 2.**
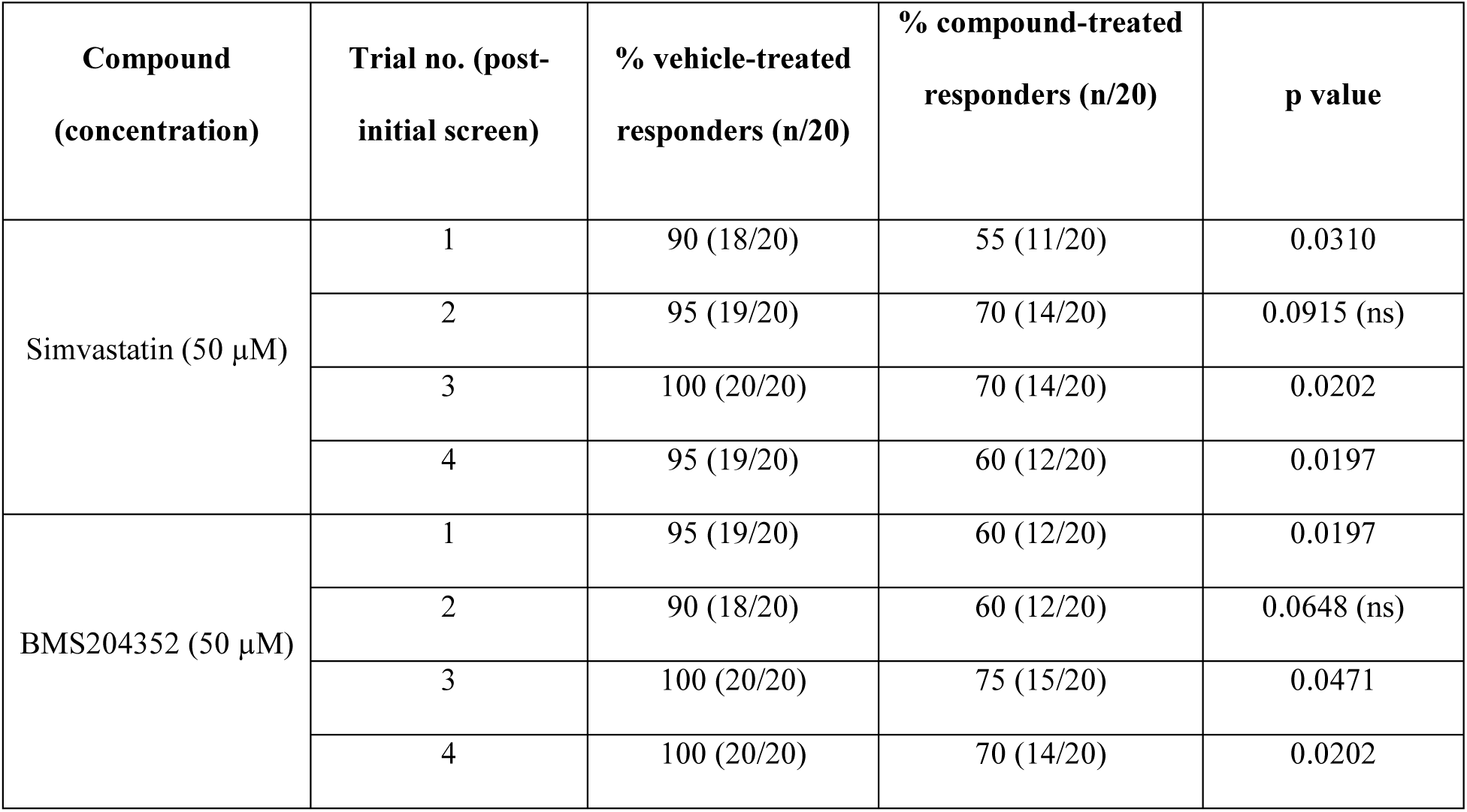
Outcomes of four independent trials to validate the efficacy of simvastatin and BMS-204352 following the initial compound screen. The data in the table refer to that depicted in Figure 4A-B. For each trial, the number of vehicle- and compound-treated *Nf1*^*P1*^ larvae responding to the stimulus is given, both as a fraction of total larvae tested (*n* = 20 per trial) and as a percentage. The outcome (p value) of the statistical comparison between these two groups in each trial is also stated. Both compounds reduced the number of responding larvae in all four validation trials, with this reduction reaching significance (p<0.05) in 3 out of 4 trials for each. For trials in which the reduction was not significant, p values were nevertheless approaching 0.05. Statistical analysis (via Fisher’s exact text) was carried out on raw data rather than percentage values. ns = not significant.

| Compound<br>(concentration) | Trial no. (post-<br>initial screen) | % vehicle-treated<br>responders (n/20) | % compound-treated<br>responders (n/20) | p value |
| --- | --- | --- | --- | --- |
| Simvastatin (50 μM) | 1 | 90 (18/20) | 55 (11/20) | 0.0310 |
|  | 2 | 95 (19/20) | 70 (14/20) | 0.0915 (ns) |
|  | 3 | 100 (20/20) | 70 (14/20) | 0.0202 |
|  | 4 | 95 (19/20) | 60 (12/20) | 0.0197 |
| BMS204352 (50 μM) | 1 | 95 (19/20) | 60 (12/20) | 0.0197 |
|  | 2 | 90 (18/20) | 60 (12/20) | 0.0648 (ns) |
|  | 3 | 100 (20/20) | 75 (15/20) | 0.0471 |
|  | 4 | 100 (20/20) | 70 (14/20) | 0.0202 |

**Figure 3.**
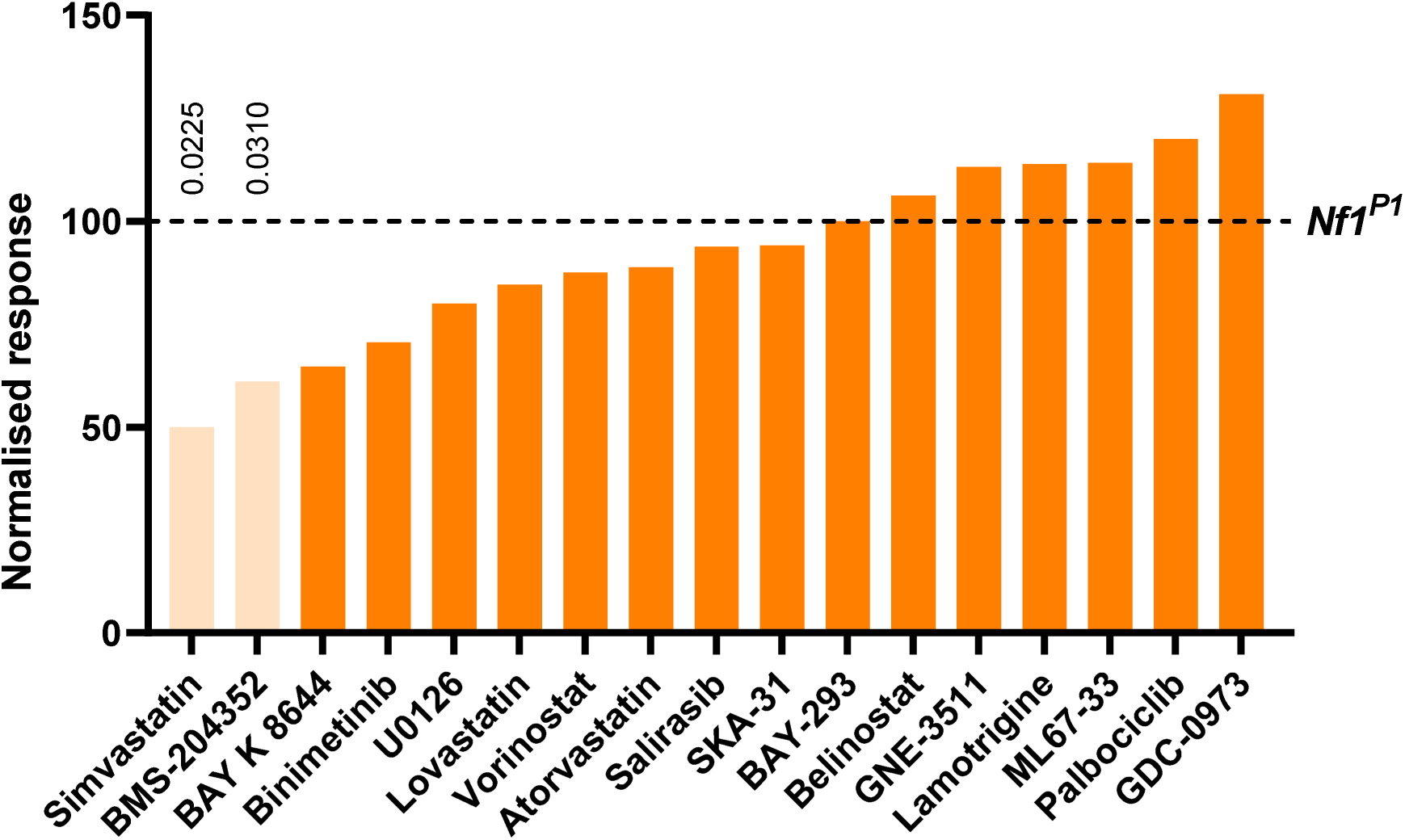
A targeted pharmacological screen identifies simvastatin and BMS-204352 to improve tactile hypersensitivity in *Nf1*^*P1*^ larvae. Twenty compounds were screened for their ability to reduce the number of *Nf1*^*P1*^ larvae responding to a mechanical stimulus, with three proving toxic at the concentration (50 µM) tested. Only administration of simvastatin (50% of vehicle-treated *Nf1*^*P1*^) and BMS-204352 (61.1% of vehicle-treated *Nf1*^*P1*^) resulted in a significant decrease in the number of responding larvae. Data are presented as a percentage of the number of *Nf1*^*P1*^ larvae raised on an equivalent concentration of DMSO (vehicle) that responded to the stimulus (dotted line = 100%). Statistical comparisons were carried out on raw data using Fisher’s exact test between compound- and vehicle-treated *Nf1*^*P1*^ larvae.

**Figure 4.**
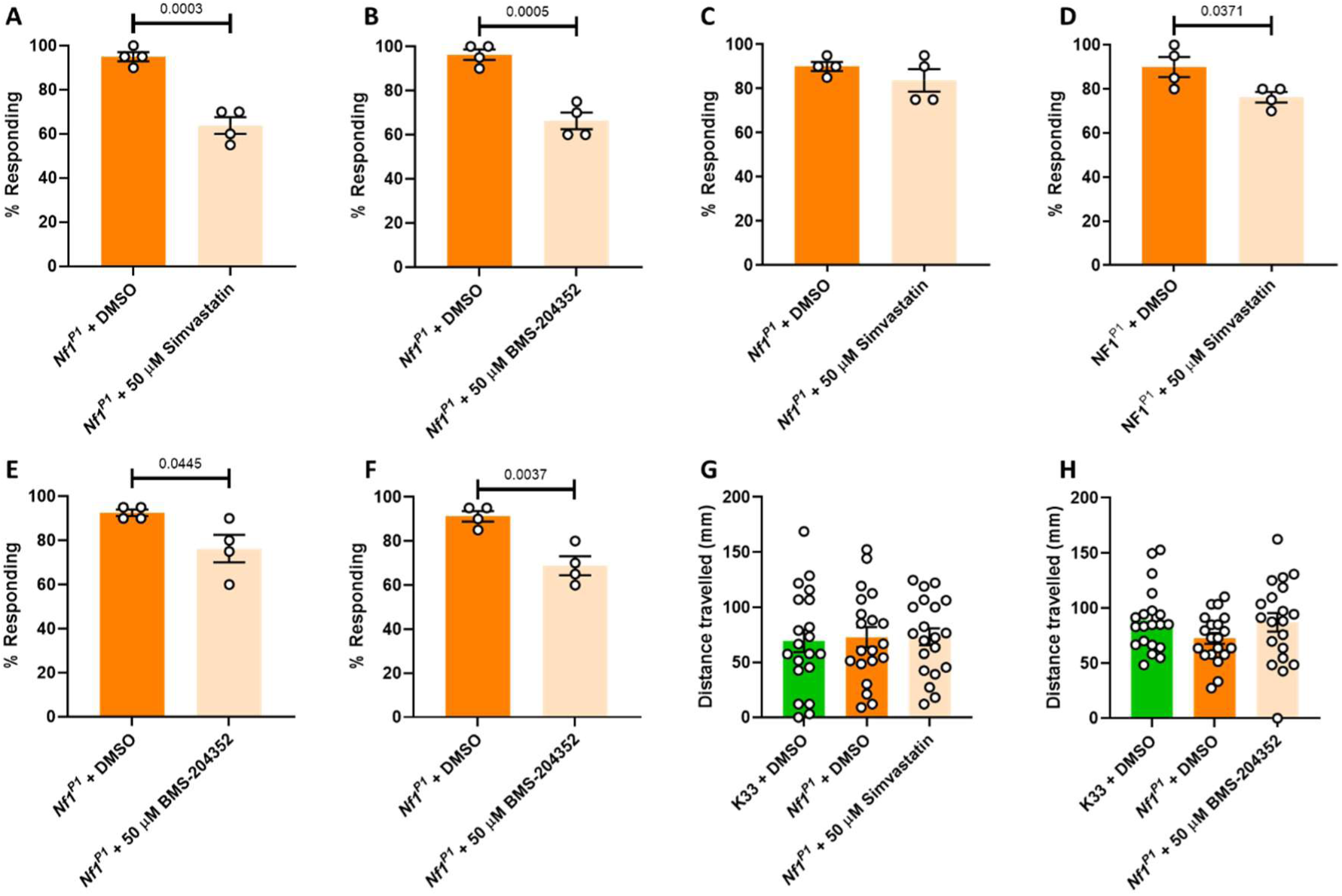
Simvastatin and BMS-204352 consistently improve, but do not fully rescue, tactile hypersensitivity in *Nf1*^*P1*^ larvae, while having no effect on overall activity. *A)* Across 4 independent trials, 50 µM Simvastatin significantly reduces the mean percentage responding larvae, as does *B)* 50 µM BMS-204352. *C)* The presence of simvastatin solely during the embryo has no impact on the percentage of *NF1*^*P1*^ larvae responding to stimulation (p=0.3026). *D)* Conversely, exposing only larval progeny to simvastatin results in a significant reduction in the mean percentage of responding *Nf1*^*P1*^ larvae. *E)* The presence of BMS-204352 solely during the embryo results in a significant reduction in the percentage of *Nf1*^*P1*^ larvae responding to stimulation. *F)* The percentage of responding *Nf1*^*P1*^ larvae is also significantly reduced when only larval progeny are exposed to BMS-204352. *G)* K33 and *Nf1*^*P1*^ larvae do not significantly differ in their total distance travelled over a 3-minute period at room temperature, nor is this impacted by treatment with 50 µM simvastatin (p=0.9518). *H)* 50 µM BMS-204352 treatment also does not alter distance travelled in *Nf1*^*P1*^ larvae, which again show no significant difference in crawling behaviour compared to K33 controls (p=0.1881). In panels A-B, each data point represents the percentage responding larvae from a single trial, with *n* = 20 per trial, such that *n* = 80 larvae overall. Data in these panels were analysed via unpaired, two-tailed student’s *t*-test. Data in panels C-D were analysed via one-way ANOVA followed by Tukey’s post-hoc test. All data are presented as mean ± SEM.

In addition, we also tested our hypothesis that compound exposure is required in both the embryo and larval stage in order to improve sensory behaviour. Accordingly, administering simvastatin solely during the embryo had no significant effect on the number of responding *NF1*^*P1*^ larvae (Figure 4C). By contrast, raising larvae on simvastatin, without embryonic exposure, significantly reduced the likelihood of a nocifensive response (Figure 4D), but to a lesser degree than when exposing both embryo and the larva (Figure 4A). These observations are consistent with our assumption that, based on *Nf1* knockdown data (Figure 1), a compound must be present during both embryonic and larval stages if it is to exert its optimal effect. However, in contrast to simvastatin, BMS-204352 administration during the embryo or larvae alone was sufficient to significantly reduce the mean percentage of responding larvae (Figure 4E-F), although the effect was stronger with larval exposure. Indeed, administering BMS-204352 solely to larvae appears to be similarly effective as administering the drug throughout both stages (Figure 4B), with changes from 91.3 ± 2.4 % to 68.8 ± 4.3 % responding (Figure 4F), and 96.3 ± 2.4 % to 66.3 ± 3.8 % responding (Figure 4A), respectively. Possibly, targeting a mechanism that may occur downstream of excessive Ras signalling, such as disrupted BK_Ca_ activity, is a more robust way of improving behaviour, such that pharmacological manipulation is still feasible in the larva.

While additional concentrations (10, 25, and 100 µM) of the two hit compounds were also tested, we found 50 µM the most consistently effective (data not shown). Furthermore, we also considered the possibility that one or both compounds reduce the likelihood of a nocifensive response in *Nf1*^*P1*^ larvae via a non-specific, sedative effect. To test this, the impact of each compound (50 µM) on larval mobility was measured over a 3-minute period. We observed no significant difference in the mean distance travelled by *Nf1*^*P1*^ and K33 larvae (Figure 4G-H), nor was distance influenced in *Nf1*^*P1*^ larvae by either simvastatin (Figure 4G) or BMS-204352 (Figure 4H). This supports the effect of these compounds being specific to tactile hypersensitivity, and not due to a global reduction in activity.

### Simvastatin and BMS-204352 improve synaptic transmission deficits at the *NF1*^*P1*^ larval NMJ

Tactile hypersensitivity in *Nf1*^*-/-*^ larvae is associated with impaired synaptic transmission at the NMJ, with both phenotypes arising in a Ras-dependent manner (Dyson et al., 2022). Therefore, we next investigated the effect of simvastatin (Figure 5A-F) and BMS-204352 (Figure 5G-L) exposure throughout embryonic and larval stages on activity at this peripheral glutamatergic synapse. Exposure to simvastatin (50 µM) led to a significant reduction in the typically enhanced mEJP frequency of *Nf1*^*P1*^ larvae, although rescue was only partial since this was still significantly greater than that of vehicle-treated K33 controls (Figure 5B). Conversely, the same treatment led to full rescue of the increased quantal size (mEJP amplitude) and reduced quantal content (Figure 5C-D). There was no effect on EJP amplitude, which is also unaltered by the *Nf1*^*P1*^ mutation (Figure 5A), and simvastatin treatment had no effect on any parameter measured in K33 larvae. Similar effects on EJP amplitude (Figure 5G), mEJP frequency (Figure 5H), and mEJP amplitude (Figure 5I) were also observed following BMS-204532 (50 µM) treatment, with the exception that the increase in quantal content in *Nf1*^*P1*^ larvae treated with BMS-204352, compared to those treated with DMSO, was not significant (Figure 5J). However, the quantal content of *Nf1*^*P1*^ larvae treated with BMS-204352 was also not significantly different from that of vehicle-treated K33 larvae. Again, there was no effect of BMS-240352 on any parameter measured in K33 larvae.

**Figure 5.**
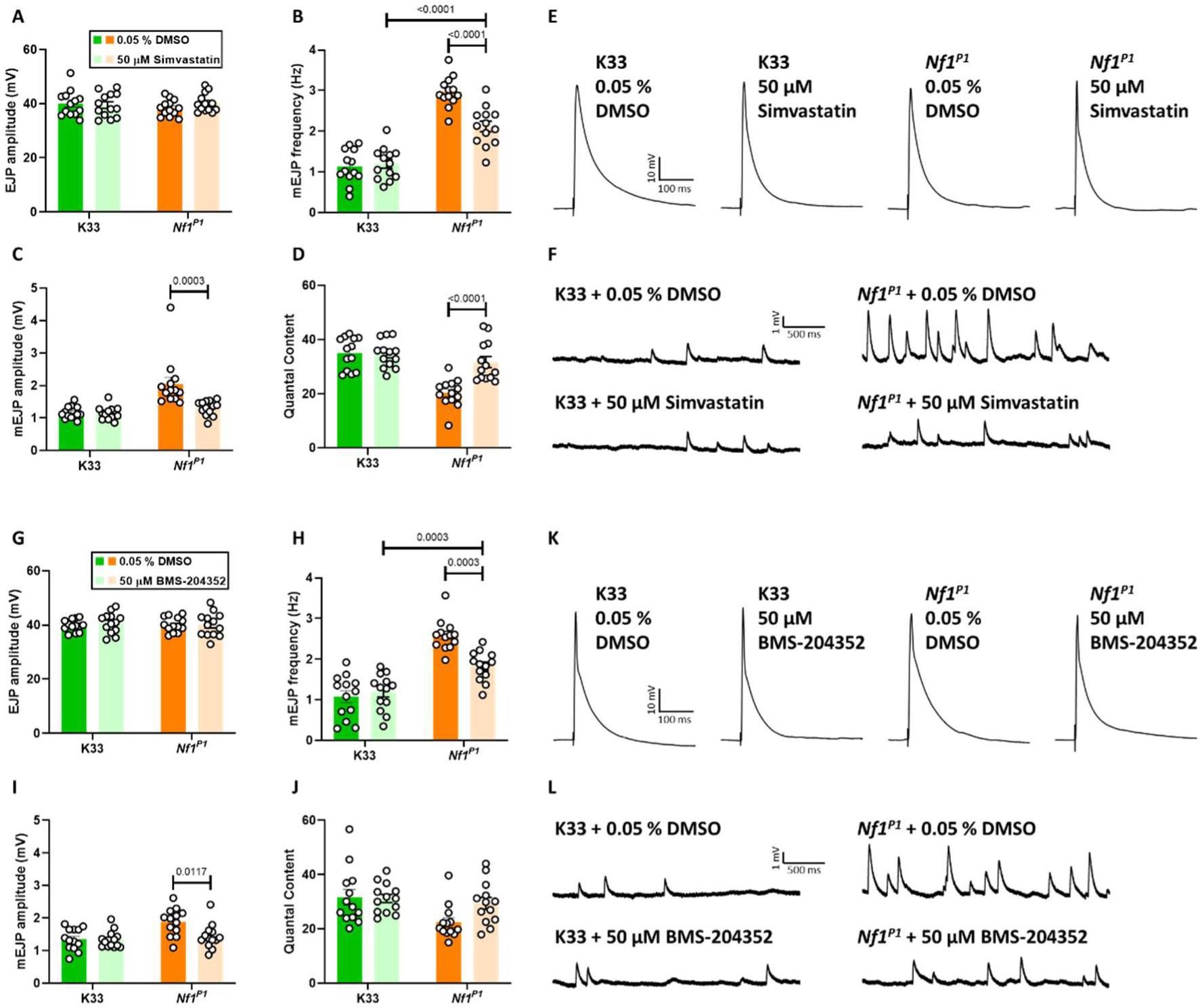
Simvastatin and BMS-204352 improve synaptic transmission deficits at the *Nf1*^*P1*^ larval NMJ, and have no impact upon normal transmission in K33 larvae. *A) S*imvastatin (50 µM) treatment has no effect on EJP amplitude in either *Nf1*^*P1*^ or K33 larvae (p=0.2793). *B)* The increased mEJP frequency of *Nf1*^*P1*^ larvae is reduced following simvastatin treatment, although this is still significantly greater than that of vehicle-treated K33 larvae. There is no significant difference between vehicle- and simvastatin-treated K33 larvae (p=0.9784). *C)* Simvastatin rescues the enhanced mEJP amplitude of *Nf1*^*P1*^ larvae to values indistinguishable from those of vehicle-treated K33 larvae (p=0.8097), which do not show a significant difference compared to simvastatin-treated K33 larvae (p>0.9999). *D)* Simvastatin rescues the reduced quantal content of *Nf1*^*P1*^ larvae to values indistinguishable from those of vehicle-treated K33 larvae (p=0.4934), which also do not show a significant difference compared to simvastatin-treated K33 larvae (p=0.9989). *E-F)* Representative traces of data presented in panels A-D. *G)* BMS-204352 (50 µM) treatment has no effect on EJP amplitude in either *Nf1*^*P1*^ or K33 larvae (p=0.5878). *H)* The increase in mEJP frequency of *Nf1*^*P1*^ larvae is reduced following BMS-204352 treatment, although this is still significantly greater than that of vehicle-treated K33 larvae. There is no significant difference between vehicle- and simvastatin-treated K33 larvae (p=0.9098). *I)* BMS-204352 rescues the enhanced Mejp amplitude of *Nf1*^*P1*^ larvae to values indistinguishable from those of vehicle-treated K33 larvae (p=0.9211), which do not show a significant difference compared to BMS-204352-treated K33 larvae (p=0.9996). *J)* The increase in quantal content in *Nf1*^*P1*^ larvae following BMS-204352 treatment is not significant (p=0.1075), however, the quantal content of BMS-204352 treated *Nf1*^*P1*^ larvae also does not differ from that of vehicle-treated K33 larvae (p=0.8849). BMS-204352 does not significantly alter quantal content in K33 larvae either (p=0.9991). All statistical comparisons were made via two-way ANOVA followed by Tukey’s post-hoc test, in which each genotype + treatment group was compared to all others. *n* = 13 for each group. All data are presented as mean ± SEM. Although not explicitly stated in the figure or legend, in panels B-D and H-J, mEJP frequency and amplitude were both significantly increased, and quantal content significantly reduced, in vehicle-treated *Nf1*^*P1*^ larvae relative to vehicle-treated K33 controls, as would be expected in larvae lacking *Nf1* expression (Dyson et al., 2022).

## Discussion

Clinical trials have thus far failed to identify treatments that effectively and consistently help to manage core ASD symptomatology, including abnormal responses to sensory stimuli. To address this, we examined whether compounds that target either Ras/MAPK signaling or ion channel activity can improve tactile hypersensitivity in *Nf1*^*P1*^ larvae. We also investigated the temporal requirements for *Nf1* in regulating the two ASD-associated phenotypes employed in this study. This is because the onset of certain ASD symptoms may occur as a result of specific developmental disruption; consequently, the time during which treatment is administered may be of equal importance as the molecular mechanism targeted. For example, in a mouse model of the syndromic ASD Tuberous Sclerosis, a 4-week regimen of rapamycin treatment beginning at postnatal day 7 prevented the occurrence of impaired social behaviour, even four weeks after the end of treatment (Gibson et al., 2022). Conversely, rapamycin administration starting at 10 weeks of age was unable to rescue the deficit (Tsai et al., 2018). Here, we find that *Nf1* likely regulates NMJ synaptic transmission via a developmental role, since its early, embryonic, downregulation is sufficient to induce persistent deficits in the third instar larva. This is not necessarily surprising, given that the loss of function of other ASD-associated genes required for NMJ development, such as the adaptor protein *ank2* (Koch et al., 2008; Stevens and Rasband, 2021) and the presynaptic adhesion molecular *neurexin-1* (Li et al., 2007; Hu et al., 2019), has also been shown to disrupt third instar larval NMJ synaptic transmission. However, in GABAergic hippocampal neurons of mice, loss of *Nf1* causes enhanced inhibitory transmission via increased Ras/MAPK-dependent phosphorylation of synapsin I, a synaptic vesicle protein that, upon phosphorylation, dissociates from vesicles to facilitate their recruitment to the readily-releasable pool (Cui et al., 2008). Thus, the mechanism of *Nf1*-dependent transmission here appears to be physiological, rather than developmental. Yet, differences in how the loss of *Nf1* impacts neurotransmission in the mouse hippocampus relative to that at the *Drosophila* NMJ have already been discussed (Dyson et al., 2022), suggesting that *NF1* may regulate this process via multiple mechanisms, and/or in a cell type-dependent manner. It should also be noted that, given the temporal resolution of the UAS/GAL4/GAL80^ts^ system (McGuire et al., 2004), we cannot entirely rule out an additional requirement for *Nf1* in the early first instar larval stage.

In contrast to NMJ phenotypes, constitutive loss of *Nf1* is required to induce larval tactile hypersensitivity. One possible explanation for this difference is that, while abnormal synaptic transmission may contribute to tactile hypersensitivity, other pathophysiological mechanisms, occurring within the larval stage are also necessary, such that the former is insufficient to induce the behavioural phenotype alone. Alternatively, whilst *Nf1* may regulate synaptic transmission at the NMJ in a developmental manner, it may not function similarly at central synapses, as it was previously shown that tactile hypersensitivity in *Nf1*^*P1*^ larvae likely arises from a central, cholinergic deficit (Dyson et al., 2022). It is also conceivable that *Nf1* may be required for some later compensatory mechanism that does not directly correct synaptic dysfunction but, nevertheless, ensures that it does not lead to changes in mechanosensory behaviour. This may mean that, in the *Nf1*^*P1*^ mutant, no compensation occurs following early changes to synaptic transmission because the *Nf1* gene is permanently deleted; conversely, in the UAS/GAL4/GAL80^ts^ paradigm, re-expression of *Nf1* in later larval stages following embryonic knockdown permits compensatory changes to prevent deficits in behaviour. Future work is required to investigate these, and other, possibilities, and to also narrow down the *Nf1*-dependent critical period of synaptic transmission.

Regardless of the precise timings of *Nf1* function, we have identified two compounds that are able to improve tactile hypersensitivity and impaired synaptic transmission in *Nf1*^*P1*^ larvae. These are the HMG CoA-reductase inhibitor simvastatin, and the BK_Ca_ channel activator BMS-204352. Simvastatin, currently prescribed as a cholesterol-lowering medication, has been examined in clinical trials for its efficacy in treating behavioural symptoms of Neurofibromatosis Type 1, with mixed success. It did not improve cognition in two large randomized controlled trials of children and adolescents (Krab et al., 2008; van der Vaart et al., 2013), possibly because the intervention occurred too late, with it being suggested that simvastatin treatment may have been beneficial in younger children (van der Vaart et al., 2013). Our data support this, since the improvement in tactile hypersensitivity in *NF1*^*P1*^ larvae was strongest when administration of the compound throughout the larval stage was combined with embryonic exposure. Indeed, some improvement in ASD symptomatology following simvastatin treatment was demonstrated in a more recent, pilot study of younger children with Neurofibromatosis Type 1 (Stivaros et al., 2018). However, if a drug must indeed be present during early (e.g. embryonic) as well as later development to elicit an effect, as our data suggest, this may entail considerable practical and ethical implications in the clinic.

BMS-204352 currently has no clinical application, having originally been developed as a therapy for acute ischemic stroke but displaying no benefit over placebo in phase III trials (Jensen, 2002). Its efficacy here may implicate the reduced expression, or impaired activity, of BK_Ca_ channels in the mechanism giving rise to tactile hypersensitivity and impaired transmission in *Nf1*^*P1*^ larvae. Roles for *Nf1* in regulating potassium currents have been demonstrated previously, as the post-synaptic K^+^ current typically elicited by application of PACAP38 is diminished at the *Nf1*^*P1*^ NMJ (Guo et al., 1997), and treatment with apamin, an inhibitor of the SK_Ca_ channel, rescues spatial learning deficits in *Nf1*^*+/-*^ mice (Kallarackal et al., 2013). Furthermore, haploinsufficiency of *KCNMA1* (encoding BK_Ca_) has been previously identified in an ASD individual, resulting in a diminished BK_Ca_ current in patient-derived lymphoblastoid cells that was enhanced via application of BMS-204352 (Laumonnier et al., 2006). BMS-204352 treatment has also been shown to rescue sensory hypersensitivity (Zhang et al., 2014), altered social preference (Hebert et al., 2014), and hyperactivity in an unfamiliar environment (Carreno-Munoz et al., 2018) in mouse models of the syndromic ASD Fragile X. In addition, BMS-204352 and an alternative BK_Ca_ activator, BMS-191001, were found to rescue structural deficits (Hebert et al., 2014) and hyperexcitability (Zhang et al., 2014), respectively, in dendrites. Thus, our finding may hint at a shared mechanism underlying sensory hypersensitivity, and potentially other symptoms, in different forms of syndromic ASD.

Despite demonstrating the consistent efficacy of these compounds in improving *NF1*-dependent tactile hypersensitivity in four subsequent trials, we were unable to fully rescue the phenotype, since K33 control larvae typically show a 0 – 20% likelihood of responding in the mechanoreception assay (Dyson et al., 2022). One explanation is that we did not test compounds at optimum concentrations. Alternatively, it may be because the target of the drug is not the only mechanism concerned. For instance, while BK_Ca_ dysfunction may contribute to tactile hypersensitivity, the dysfunction of other ion channels, for example, may also be involved, which must also be pharmacologically corrected to completely rescue the phenotype. However, it is difficult to apply this reasoning to simvastatin: although we have not shown biochemically that simvastatin functions in our assay by targeting Ras, the most parsimonious explanation for its effect is that, by inhibiting HMG CoA-reductase, it prevents the farnesylation of Ras and its subsequent anchoring to the plasma membrane (Li et al., 2005). Yet, genetically attenuating Ras protein expression via RNA interference fully rescues the hypersensitivity phenotype (Dyson et al., 2022).

In summary, we have shed light on the temporal requirements for the NF1 protein in regulating two previously identified larval phenotypes that arguably parallel ASD symptomatology and aetiology. Subsequently, we identified two compounds that improve, but do not necessarily fully rescue, the same phenotypes. Despite the previous failure of simvastatin to improve cognitive symptoms in children with Neurofibromatosis Type 1 (Krab et al., 2008; van der Vaart et al., 2013), our findings suggest that the further investigation of this compound, and of BMS-204352, specifically in the context of ASD may be warranted.

## Notes

Authors report no conflict of interest.

This work was funded by a Medical Research Council Doctoral Training Partnership to A.D., and by funding from the Biotechnology and Biological Sciences Research Council (BB/L027690/1) to R.A.B. S.G. is supported by the Neurofibromatosis Therapeutic Acceleration Program (NTAP) Francis Collins Scholarship. D.G.E. is supported by the Manchester NIHR Biomedical Research Centre (IS-BRC-1215-20007).

### Competing Interest Statement

The authors have declared no competing interest.

